# Computationally Designed Carbonic Anhydrase Inhibitors Derived from Sulfonamides for Treatment of Glaucoma

**DOI:** 10.1101/2023.02.25.529947

**Authors:** Danielle J. Caruso, Morgan P. Connolly, Tram Q. Nguyen, Justin B. Siegel

## Abstract

Glaucoma is a known contributor to blindness in adults. This disease occurs as a result of overactivity of the carbonic anhydrase II (CAII) enzyme. Several of the commonly used sulfonamide drugs have been used to treat the symptoms of glaucoma but have been found to have low potency. To develop a more effective drug for the treatment of glaucoma, two novel CAII inhibitor drugs were designed through bioisoteric replacement and chemical intuition. This research discusses the development of these CAII inhibitor drugs from the existing glaucoma drug known as dichlorphenamide. The drugs created in this research were found to have a better docking score within the CAII binding site than that of dichlorphenamide. The drugs proposed in this paper would need to undergo further research to determine laboratory synthesis and potential clinical trialing.

## Introduction

Glaucoma is one of the main causes of blindness in adults over the age of 60 years old.^1^ This disease of the eyes is caused by an increase in intraocular pressure due to an overproduction of aqueous humor liquid. Regulation of many fluids throughout the human body is attributed to the carbonic anhydrase II (CAII) enzyme which functions in the red blood cells, pancreatic cells, renal tubules, retina and lens.^1^ This enzyme is responsible for catalyzing the conversion of carbon dioxide and water into bicarbonate and a proton. CAII is available at high concentrations in the cellular sites of the eyes, thus allowing its production of bicarbonate ions to regulate fluid in the eyes. When there is overactivity of the carbonic anhydrase enzyme, an overproduction of fluid in the eyes can occur. This can lead to an increase in pressure within the eye that often causes nerve damage or glaucoma when untreated.^2,3^

To reduce the fluid levels in the eye to avoid glaucoma, carbonic anhydrase inhibitor drugs have been studied as a form of treatment. Recently, the well-known sulfonamide group of drugs have been found to be effective in their treatment of glaucoma.^4^ Sulfonamides, also known as sulfa drugs, are common antibacterial drugs that have been used to treat various infections and are also currently being studied for their effectiveness in anti-tumor activity.^1^ Among the common sulfonamides used specifically to treat glaucoma are acetazolamide, methazolamide, ethoxozolamide, and dichlorphenamide (Figure 1). These drugs have been found to exhibit promising carbonic anhydrase inhibition, but also tend to have low potency and adverse effects to the liver and kidney if taken for a prolonged period.^1^

**Figure 1:**
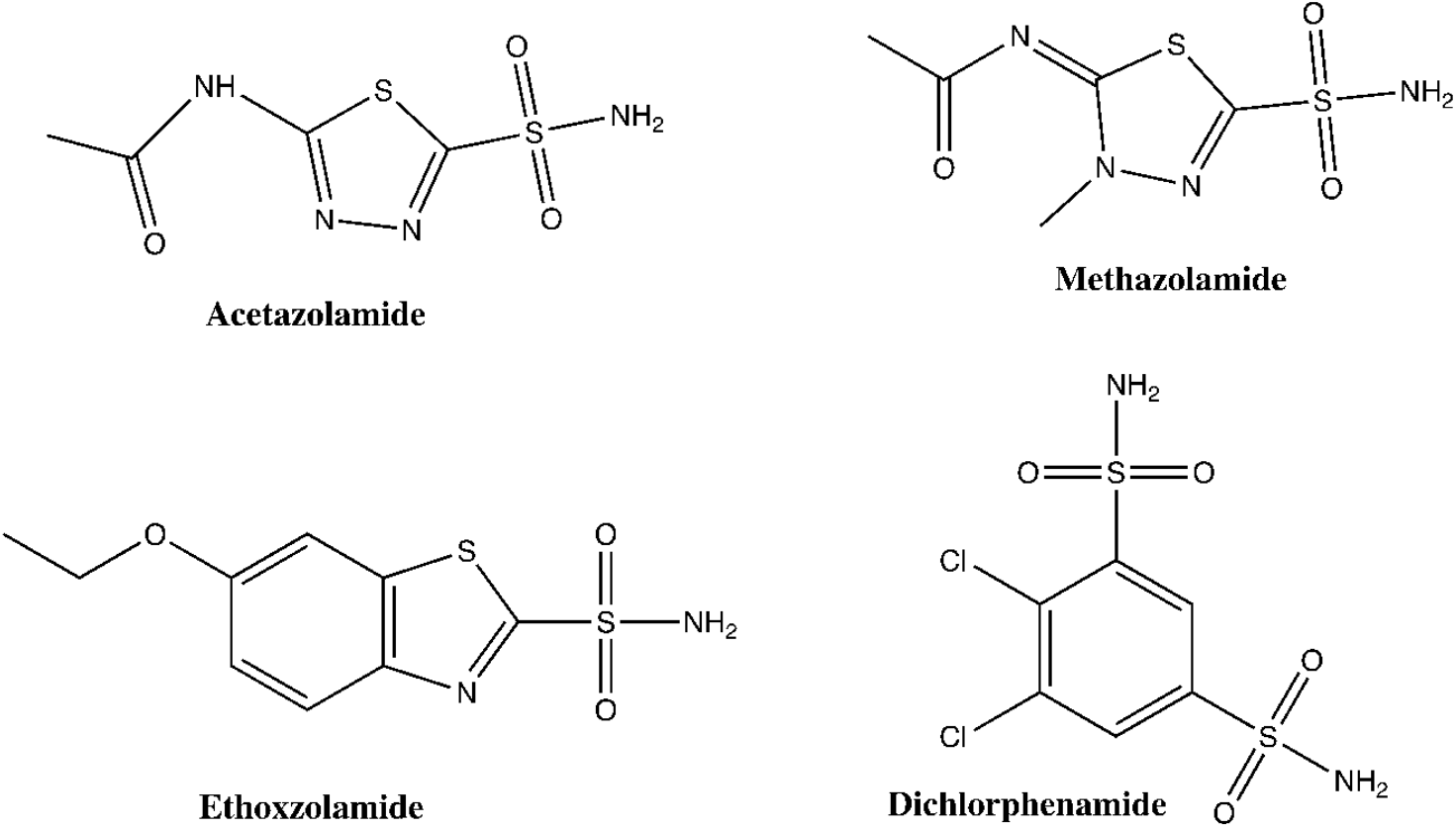
2D chemical structures of acetazolamide, methazolamide, ethoxzolamide, and dichlorphenamide.

The active site of carbonic anhydrase contains a Zn^2+^ cation that interacts with His 119, His 96, and His 94 (Figure 2). This Zn^2+^ cation interacts with nearby water molecules to initiate the interconversion of carbon dioxide to bicarbonate. However, when the sulfonamide inhibitors interact with carbonic anhydrase II, the histidine residues that previously interacted with Zn^2+^ are now bound to the inhibitor drug. The inhibition of CAII ultimately leads to the reduction in aqueous humor production in the eyes, thus reducing the intraocular pressure to reduce the disease state.^3,4^

**Figure 2:**
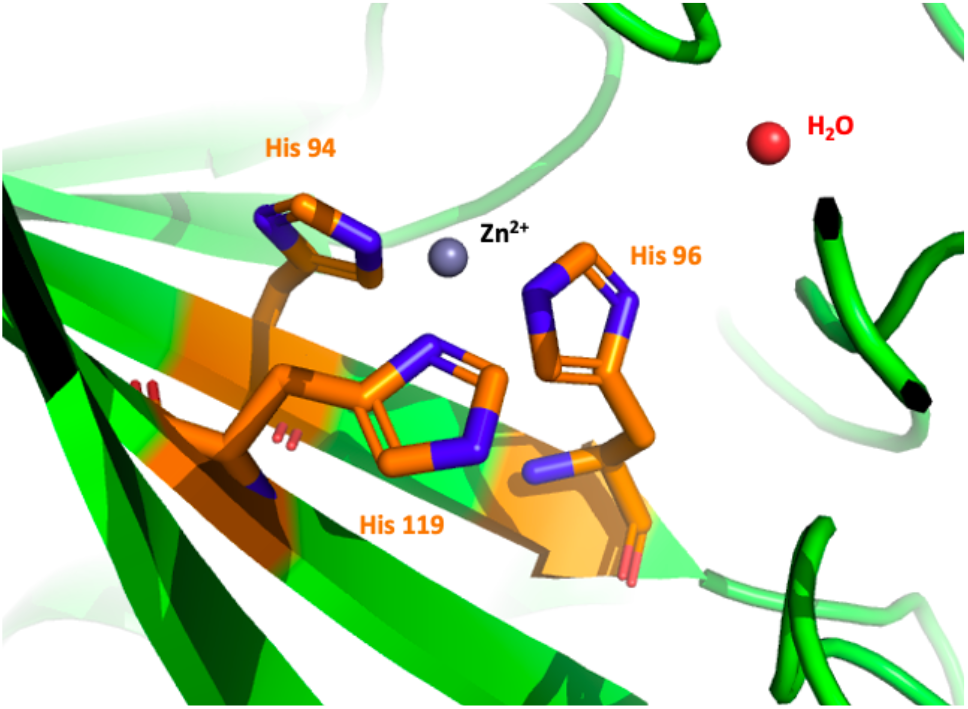
The active site of carbonic anhydrase containing the zinc ion and water molecule that interact for formation of bicarbonate.

Computational docking simulations of the four common sulfonamides used for the treatment of glaucoma showed that acetazolamide, methazolamide, and ethoxozolamide present higher binding affinity in the CAII active site than that of dichlorphenamide. Based on docking results, we proposed that the low binding affinity for dichlorphenamide may be due to its inability to effectively fit in the binding pocket of CAII as well as the other drugs. This research discusses the use of computational drug design and screening methods to optimize the structure of dichlorphenamide to develop sulfa drug candidates with improved binding.

## Methods

The crystal structure for human carbonic anhydrase II in complex with the natural ligand carbon dioxide (5YUI) was obtained through RCSB Protein Data Bank.^9^ The 5YUI crystal structure was uploaded to PyMOL to visualize interactions between the amino acid residues of CAII and the docked drug candidates. Crystal structures of CAII in complex with sulfonamides (3BL1) and acetazolamide (3HS4) were also used to visualize key interactions for inhibition but were not used for the drug docking process.^10,11^

Gaussview and Gaussian 90W software were used for 3D visualization and semi-empirical optimization of the drug candidates. Conformation libraries were generated using OMEGA and analyzed in VIDA. Using FRED and MakeReceptor, docking scores for the drugs within the CAII active site were generated ^13^ vBrood 2.0 was used to create drug candidates through bioisosteric replacement of functional groups on the dichlorphenamide drug. The ADMET properties of the drugs, including bioavailability, potential aggregation, toxicophoric groups, and Lipinski’s Rule of 5, were generated using the OpenEye Filter.

Protein BLAST from NCBI was utilized to identify species with highly homologous proteins to human CAII.^14^ The protein sequence of CAII in *Mus musculus* was displayed in Jalview software to view the conserved catalytic residues to provide insight into pre-clinical trial opportunities.

## Results and Discussion

### Analysis of Known Carbonic Anhydrase Inhibitors

Four sulfonamide carbonic anhydrase inhibitor drugs; acetazolamide, methazolamide, ethoxozolamide, and dichlorphenamide; were analyzed and used as the basis for design of new drug candidates. The properties of each of these molecules shows them to be successful inhibitor drugs for CAII, however, they tend to lack potency.^1^ After each of the drugs were docked in the CAII active site using FRED, docking scores were used to assess the drugs’ interactions with the active site.^13^ The FRED Chemgauss 4 docking scores obtained for acetazolamide, methazolamide, ethoxozolamide, and dichlorphenamide were -6.24, -5.90, -5.50, and -4.16 respectively (Table 1). Analogs of the drug dichlorphenamide were developed as opposed to the other three drugs due to the dichlorphenamide possessing the weakest docking score.

**Table 1:**
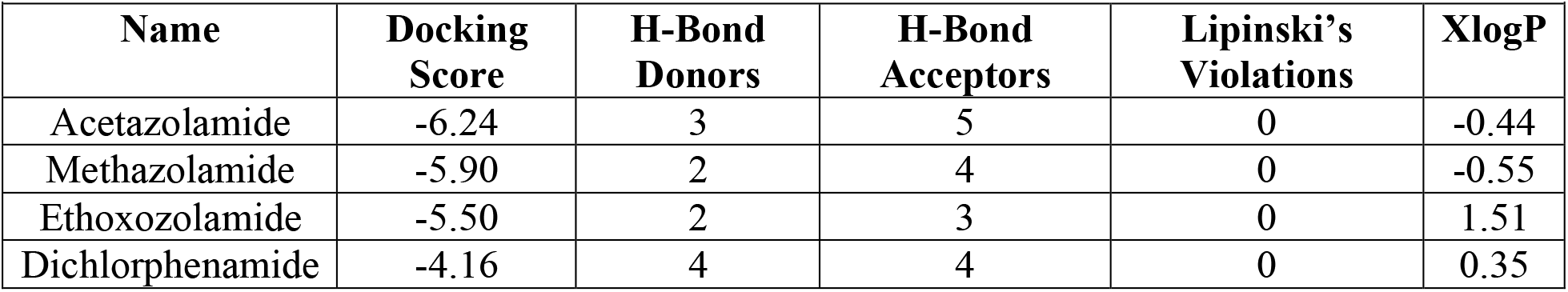
FRED docking scores and ADMET properties of four known CAII inhibitors.

In addition to docking, the absorption, distribution, metabolism, excretion, and toxicity (ADMET) properties of known CAII inhibitors were investigated. The results of this analysis, as given by the OpenEye Filter, showed the toxicophoric groups, Lipinski’s Rules violations, and the number of hydrogen bond donors and acceptors for the four sulfonamide drugs. None of the drugs screened violated any of Lipinski’s rules, possessed toxicophoric groups, or were identified as aggregators. However, there were variations in the number of hydrogen bonding capabilities among these compounds. Dichlorphenamide had four hydrogen bond donors and four hydrogen bond acceptors and methazolamide had four hydrogen bond acceptors and two hydrogen bond donors. Acetazolamide had five hydrogen bond acceptors and three hydrogen bond donors, whereas ethoxozolamide had three acceptors and two donors (Table 1).

Based on these analyses and the visualization of the drugs in the active site, dichlorphenamide (Figure 3) was the drug chosen as a template for modification. The shape of dichlorphenamide, as opposed to the other drugs, caused it to be unable to effectively fit into the binding pocket. This caused its docking score to be higher than that of the other drugs because it was not close enough to the histidine residues in the active site to hydrogen bond to them. Dichlorphenamide was only within hydrogen bonding distance for His 94. For an effective inhibition of the CAII enzyme, all three histidine residues involved in the Zn^2+^ complex should be inhibited. Additionally, to increase the specificity of this drug to the CAII active site, the number of interactions with active site residues should also be increased. Figure 3 shows the lack of strong interactions that dichlorphenamide makes in the CAII active site. New analogs of dichlorphenamide were created to increase its space filling in the binding pocket to promote interactions with His 119, 94, and 96.

**Figure 3:**
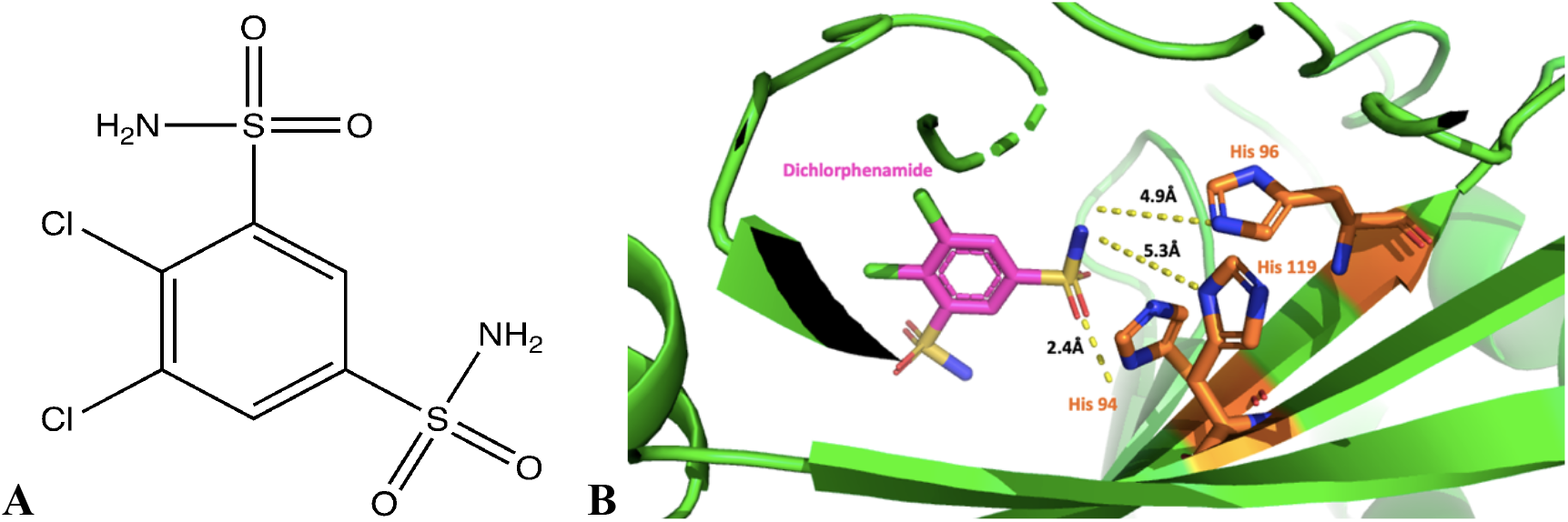
A) 2D chemical structure of dichlorphenamide. B) Dichlorphenamide (pink) docked in the active site of CAII. The interactions shown from top to bottom are between His 96 (4.9Å), His 119 (5.3Å), and His 94 (2.4Å).

### vBrood Generated Sulfonamide Drug

A computationally derived drug analog of dichlorphenamide was generated via vBrood 2.0. Dichlorphenamide was initially run through the software to by selecting the sulfone groups to undergo the bioisosteric substitution. The sulfone group was selected for alteration due to its lack of interactions in the CAII active site. Additionally, this group was selected for change because the existing shape of dichlorphenamide did not allow for binding within the enzyme active site, due to the distance between the drug and the active site amino acids being too far for viable bonding. It was proposed that selecting a larger group to replace one of the sulfone groups may shift the position of the molecule to fit better in the binding pocket. The group that was selected to replace the sulfone group was a five-membered nitrogen-containing ring with a double bonded nitrogen, an amine group, and a methyl group substituent. This group was selected due to the size and location of the amine group within the five-membered ring (Figure 4). The size of this addition allowed for a shift in the position of the molecule, ultimately allowing the amine group to hydrogen bond within the active site. Previously, in dichlorphenamide, hydrogen bonding at this location of the molecule did not exist.

**Figure 4:**
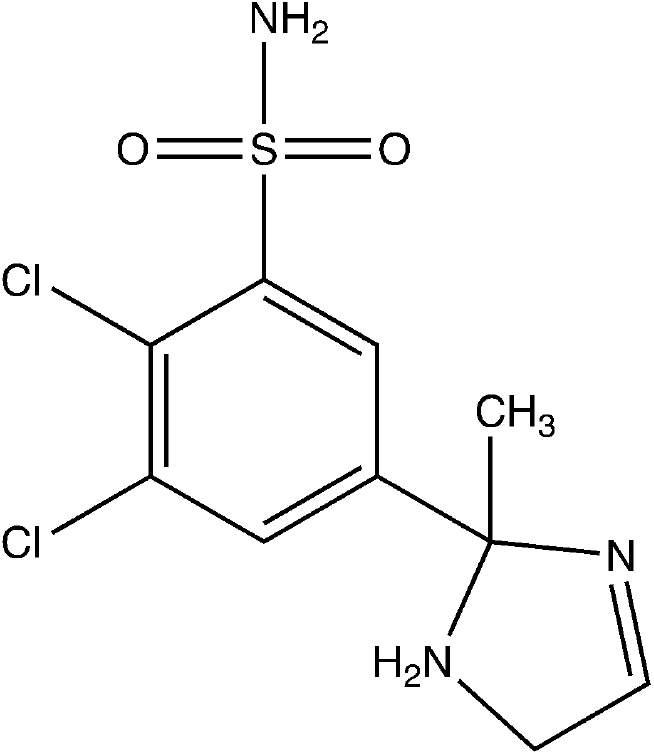
Structure of the computationally derived drug candidate (Candidate 1).

The addition of the new functional group caused the molecule to shift its position in the active site, thus allowing Candidate 1 to successfully hydrogen bond to His 94, His 96, His 119 (Figure 5). The distances to these histidine residues were shortened significantly, thus indicating stronger binding between Candidate 1 and the residues in the active site. The bond lengths for the hydrogen bond interactions between His 94, His 96, and His 119 are 2.2 Å, 2.7 Å, 2.4 Å respectively. The bond lengths presented a decrease of 0.2 Å, 2.2 Å, and 2.9 Å respectively, which brought the inhibitor close enough to both His 96 and His 119 to form a hydrogen bond. Additionally, hydrogen bond interactions between Candidate 1 and both Thr 199 and Gln 92 developed as a result of the new functional group. Gln 92 hydrogen bonds to the amine group that was added in the five-membered ring, and Thr 199 is now able to bind to the remaining sulfone group (Figure 5) The newly created interactions improved the docking score by 34.2% to -5.59. (Table 2). Although the docking score of Candidate 1 was significantly improved, Candidate 1 did possess one less hydrogen bond acceptor, potentially losing an opportunity for binding.

**Figure 5:**
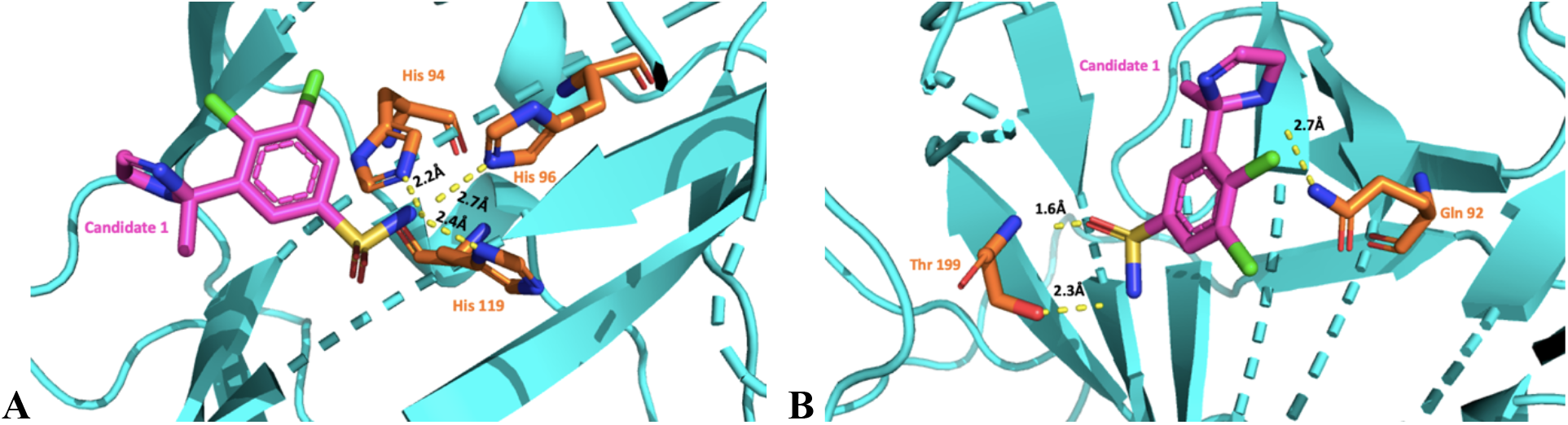
A) Interactions between Candidate 1 (pink) and the catalytic residues of His 94, His 96, and His 119. B) New hydrogen bonding interactions that Candidate 1 (pink) made with Thr 199 and Gln 92. Images generated using PyMol.

**Table 2:** Docking score and characteristics of Candidate 1.

| **Table 2:** Docking score and characteristics of Candidate 1. |  |  |  |  |  |
| --- | --- | --- | --- | --- | --- |
| **Name** | **Docking Score** | **H-Bond Donors** | **H-Bond Acceptors** | **Lipinski's Violations** | **XLogP** |
| Candidate 1 | -5.59 | 4 | 3 | 0 | 1.88 |

When Candidate 1 was run through ADMET testing, it did not display any toxicophoric or aggregator groups, nor did it violate any of Lipinski’s Rules. Because Candidate 1 did not break any of Lipinski’s rules, this drug maintains oral bioavailability. The XlogP for Candidate 1 was 1.88, as compared to the 0.35 of dichlorphenamide. This was likely due to the additional cyclic structure added. The vBrood technology proved to generate a successful candidate with an increased docking score.

### Rational Design of a Sulfonamide Drug Candidate

The next drug candidate proposed was created by structure-based rational design after analyzing where Candidate 1 could be improved. As it was discovered in Candidate 1, incorporating an additional cyclic structure in place of the pre-existing sulfone group shifted the position of the structure to allow it to have more interactions with the active site. In Candidate 2 (Figure 6), it was proposed that adding an alcohol group, as a hydrogen bond donor, to the five-membered ring in the location where there was previously a non-interacting hydrogen bond acceptor in Candidate 1, would increase the binding activity. Additionally, adding the nitrogen as a hydrogen bond acceptor to the five-membered ring in the location where there was previously a hydrogen bond donor, was proposed to enhance the chance of hydrogen bonding. This proposed structure did not have the methyl group bound to the five-membered ring, as that group did not seem to interact within the active site and displayed some strain to the molecule. This molecule was built using Gaussview and optimized through Gaussian. The docking score was determined through FRED.

**Figure 6:**
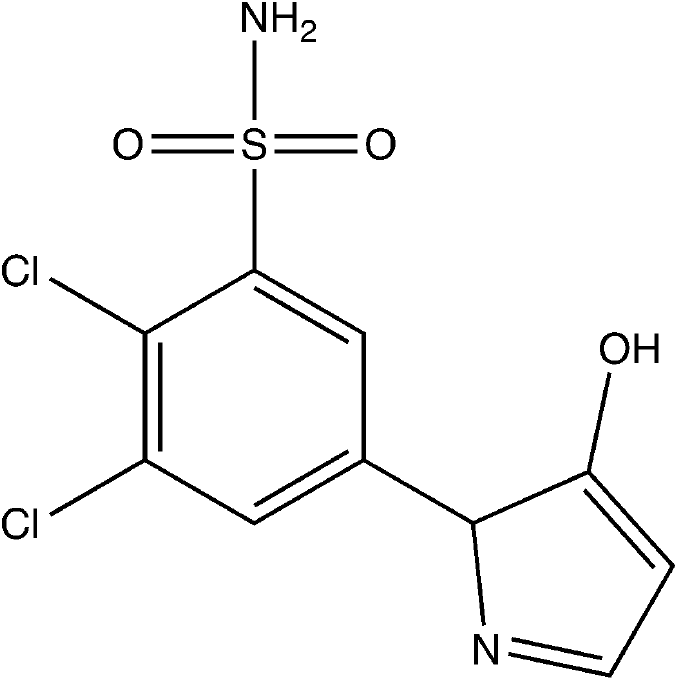
The 2D chemical structure of Candidate 2.

Candidate 2 had a docking score of -5.43, a 30.5% increase from the original -4.16 (Table 3). The improvement in docking score for Candidate 2 was likely due to the new hydrogen bonds that were made in the active site, as well as a newly available pi-pi stacking interaction with the five-membered ring (Figure 7).

**Table 3:** Docking score and characteristics of Candidate 2.

| **Table 3:** Docking score and characteristics of Candidate 2. |  |  |  |  |  |
| --- | --- | --- | --- | --- | --- |
| **Name** | **Docking Score** | **H-Bond Donors** | **H-Bond Acceptors** | **Lipinski's Violations** | **XLogP** |
| Candidate 2 | -5.43 | 3 | 3 | 0 | 1.24 |

**Figure 7:**
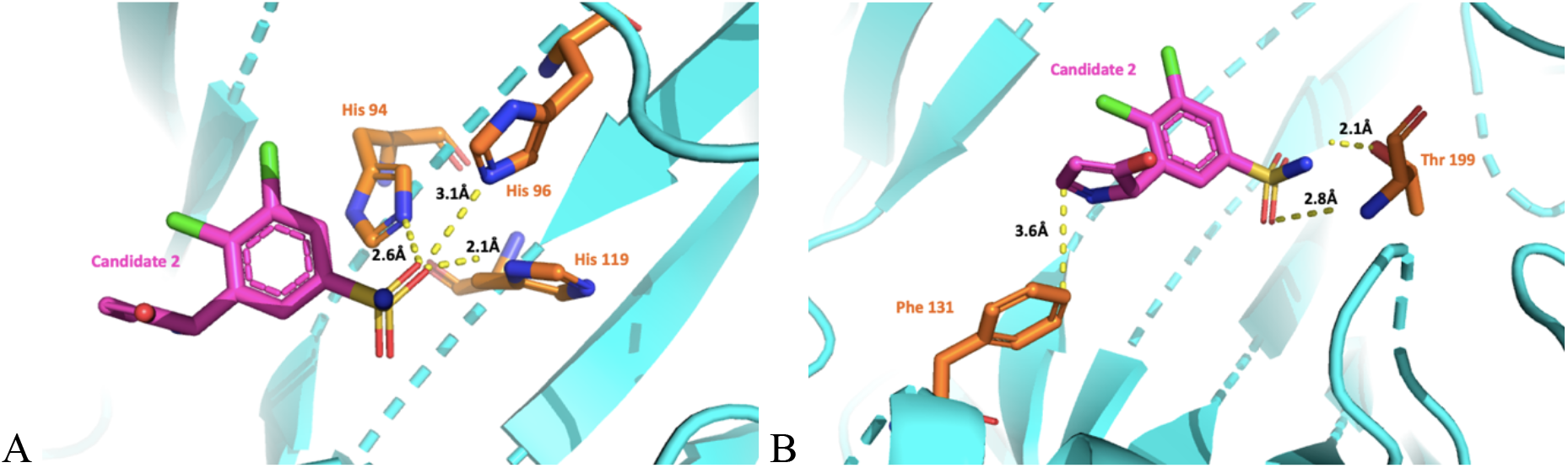
A) Interactions between Candidate 2 (pink) and His 94, His 96, and His 119 of the CAII active site. B) Interactions that were made between Candidate 2 (pink) and Thr 199 and Phe 131. Images generated using PyMol.

Similar to Candidate 1, adding in the cyclic group as a replacement for the second sulfone group is predicted to allow for a structural shift such that hydrogen bonding can occur with His 94, His 96, and His 119. The bond lengths for each of the histidine residues was shortened from the original lengths that were displayed in Figure 3. The hydrogen bond lengths for His 94, His 96, and His 119 are 2.6 Å, 3.1 Å, and 2.1 Å respectively. Aside from these hydrogen bonds contributing to the increased docking score, Candidate 2 was also able to participate in pi-pi stacking with Phe 131. Unlike in Candidate 1, the five-membered ring for Candidate 2 was shifted in a position that allowed it to lay in the same plane as Phe 131 and thus create the pi stacking interaction between the carbons in both pi systems. The bond length for this interaction was 3.6 Å. Although it was proposed that adding the alcohol group to the five-membered ring would contribute to hydrogen bonding with the nearby Pro 202 residue, this residue ended up being 4.0 Å away from the alcohol group, thus being too far to contribute to bonding. Furthermore, there were no other residues close enough to the alcohol group to participate in bonding. In future optimization, this group may be eliminated or substituted for another functional group to allow for additional binding activity. The nitrogen within the five-membered ring also did not contribute to hydrogen bonding. Though both groups were added with the intentions of hydrogen bonding capabilities, this did not occur. The pi-stacking of this five-membered ring was an unpredicted interaction.

Although the docking score for Candidate 2 was an improvement from the original drug, the fact that it has one less hydrogen bond donor and acceptor than dichlorphenamide may contribute to Candidate 2 having a higher score than Candidate 1. Despite the decreased number of hydrogen bond acceptors and donors, Candidate 2 shows promising characteristics after ADMET screening via OpenEye Filter. The XlogP for this candidate was 1.24 which is higher than the 0.35 of dichlorphenamide, but lower than the 1.88 of Candidate 1. The increase in XlogP compared to dichlorphenamide is likely due to the additional cyclic structure added. This candidate’s XlogP was lower than Candidate 1 likely because of the alcohol group that was added, causing this structure to be slightly more soluble in water. Candidate 2 does not possess any toxicophoric functional groups or aggregator properties. Additionally, this candidate does not violate any of Lipinski’s Rules, and therefore can be taken as an oral tablet as intended.

Both Candidate 1 and 2 were searched in Scifinder to determine prior use. Neither of these candidates have been proposed prior to this research. Between Candidate 1 and 2 it appears that Candidate 1 has the overall better docking score and presents more promising results.

### Prospects for Drug Candidates in Pre-Clinical Trials

Carbonic anhydrase II is a prominent enzyme in hundreds of species, but does not display the same amino acid sequence in each of these species. To determine the potential for these drug candidates to be brought to the market, an important consideration is how they would perform during pre-clinical trials. Every drug undergoes pre-clinical trials to determine the safety and efficacy of the drug before it is used in humans. Typical pre-clinical trials are performed on laboratory mice (*Mus musculus)*. For accurate and representative pre-clinical trials to take place, the species that the drugs are being tested on must have a similar amino acid sequence to the human enzyme. In the case of this study, it was discovered that the CAII enzyme is present in a homologous protein in *Mus musculus* and possesses a sequence identity of 99% to CAII of *Homo sapiens*. Having a homologous protein that functions the same way in both species is necessary for pre-clinical trialing, otherwise the drug may experience off-target effects in the other species.

To analyze the use of these drug candidates in *Mus musculus* for pre-clinical trials, the NCBI protein BLAST program along with Jalview software were used to compare the amino acid sequences present in the CAII active site in *Mus musculus* (10088) and *Homo sapiens*.^15^ For the drug candidates proposed in this study to work effectively on *Mus musculus*, the CAII residues that participate in the inhibition of CAII must be the same as they are in *Homo sapiens*. The residues that participate in the inhibition of CAII are His 94, His 96, and His 119. After visualizing the amino acid sequences through Jalview, it was determined that the *Mus musculus* CAII sequence also displays these three active site residues in the same locations as *Homo sapiens*.

Since the CAII enzyme in *Mus musculus* possesses the same key features as in *Homo sapiens*, the drug candidates could be used in pre-clinical trials in mice. It should be noted, however, that some residues differ between *Homo sapiens* and *Mus musculus* outside of the active site. Further testing would need to be done to determine if these differences could contribute to off-target effects or changes in drug performance between humans and mice. It was also discovered via UniProtKB that the CAII active site in mice is inhibited by acetazolamide, one of the sulfonamide inhibitors of CAII in *Homo sapiens*. This indicates that known sulfonamide inhibitors do effectively bind to CAII in *Mus musculus* as they would in *Homo sapiens*.^12^ The active site of CAII for *Mus musculus* with the drug candidates is not shown since the active site is highly similar to that of *Homo sapiens*.^9^

## Conclusion

Glaucoma is a serious medical condition that can lead to blindness in adults if untreated. This condition is typically managed through the means of eye drops or laser eye surgery, but there currently exists no cure for this disease. Although there are small molecule inhibitor drugs for this condition, these drugs do not display the desired potency to effectively lower the intraocular pressure.^1^ Ghorai, *et al*. (2020) proposed that the binding affinity and potency of these drugs can be improved by altering functional groups to occupy more of the binding pocket in carbonic anhydrase II. In order to work towards the improvement of these drugs, this study aimed to improve the docking scores and active site interactions of the known sulfonamide drug, dichlorphenamide.

Through computational analyses and rational design, two analogs of dichlorphenamide were proposed, Candidate 1 and 2. Both showed significant improvement in their docking scores with novel hydrogen bond interactions within the active site of CAII. These drugs possessed promising characteristics for the treatment of glaucoma, as they strongly interact with the three histidine catalytic residues. These catalytic residues, when bound to the candidates, can no longer interact with the Zn^2+^ cofactor of CAII, thus inhibiting the interconversion of carbon dioxide, and reducing the intraocular pressure of the eye.

The two candidates designed in this study were screened for ADMET properties to test for safety and oral bioavailability, however, upon metabolism the characteristics of these drugs may change. The homologous protein in *Mus musculus* was analyzed to understand this species’ potential use in pre-clinical trials to determine the effects these drugs could have in animals prior to human trials. Future research on this project may be conducted to assess the activity of these drugs *in vitro* and *in vivo* if deemed desirable.

